# Asymmetric Cell Growth and Nucleoid Displacement in *Escherichia coli*

**DOI:** 10.1101/682435

**Authors:** Manasi S. Gangan, Chaitanya A. Athale

## Abstract

Single celled growth of *Escherichia coli* is typically considered as an example symmetric division, based on the sizes of daughter cells and precision of center finding of the septum. Here, we investigate the symmetry of membrane addition in the mid-plane and DNA segregation using a video-microscopy approach. We find the membrane expansion dynamics to be asymmetric based on mid-cell photobleaching landmarks in FM4-64, used to stain the membrane. The apparent growth bias of *E. coli* does not correspond to the age of the pole. We find the membrane growth asymmetry is correlated to nucleoid displacement, consistent with ideas of coupling of cell growth and nucleoid positioning. The mobility of the actin-homolog MreB also correlates with membrane growth asymmetry, based on fluorescence recovery after photobleaching (FRAP) measurements of a YFP-fusion. These correlations suggest the small asymmetry of membrane addition observed could potentially drive nucleoid segregation and MreB mobility asymmetry in *E. coli*.

**IMPORTANCE:** Asymmetry in bacterial cell division is seen in it’s most simple form in the binomial event of protein segregation that results in ‘noise’ in equal segregation and depends on the protein copy number. In the case of specific proteins this can also affect growth, and result in differentiation. However, during the asexual division of *Escherichia coli* single cells are thought to grow symmetrically and divide equally. We find a slight but consistent asymmetry in growth based on quantitative morphometry of cell pole displacement in time-series of growing *E. coli* cells. Increased cell wall extension in one half of the cell over the other appears to explain the asymmetry in the displacement of cell poles. Interestingly the growth asymmetry is not correlated to the age of the pole (old and new). This observed asymmetry appears to correlate with the asymmetry of nucleoid segregation, resulting in the nucleoids finding the midpoints of the respective daughter cells, and formation of the septum at the geometric centre of the elongated cell just prior to division. The mobility dynamics of the cytoskeletal protein MreB, which organizes the cell membrane, is are more dynamic where membrane growth is faster. Thus we propose a linkage between this observed growth asymmetry and that of MreB dynamics.

## INTRODUCTION

Asymmetric growth in cellular systems is observed across genera in order to introduce functional diversification (Nelson, 2012). In particular, individual bacterial cells with geometrically simple shapes (e.g.: rods, ellipses, spheres) undergo symmetric cell division through asymmetry in growth and molecular localization (Kysela et al., 2013). At one extreme are bacteria such as those that belong to the group α-proteobacteria that exhibit distinct morphological asymmetry (Skerker and Laub, 2004; Brown et al., 2012), and *Caulobacter crescentus* that forms two different daughter cells- stalk and the swarmer- that have not just morphological differences, but also perform distinct functions (Siegal-Gaskins and Crosson, 2008; Skerker and Laub, 2004). While asexual division of model bacteria such as the rod-shaped gram positive *Bacillus subtilis* and gram-negative *Eschericihia coli* is symmetric, resulting in morphologically identical daughter cells, both these bacterial species can also undergo asymmetric cell division. *B. subtilis* divides asymmetrically forming a small, spore cell when faced with nutrient-poor environments (Beall and Lutkenhaus, 1991; Ben-Yehuda and Losick, 2002) while damaged proteins are selectively segregated into the ‘old’ pole during *E. coli* division, as a means of possibly improving the fitness of one of the daughter cells (Lindner et al., 2008). Unipolar and asymmetric growth of rod-shaped cells across the mid-axis of a cell has been observed in *Mycobacterium sp.* with morphological and antibiotic sensitivity differences seen in the progeny (Aldridge et al., 2011; Joyce et al., 2012).

The growth of the model bacterium, *Escherichia coli* is considered to involve dispersed regions of active growth, across the cell length, away from the poles (Kysela et al., 2013). The cytoskeletal protein MreB, an actin homolog, is essential for maintenance of the rod-shape of growing in *E. coli* cells (reviewed by Shi et al., 2018), with a minor role played by FtsZ, the tubulin homolog (Varma et al., 2007). The cell membrane growth prior to septation is driven by the synthesis of lipids and the peptidoglycan layer, with MreB anchoring some of the proteins involved in this pre-septation growth (Typas et al., 2011). However, the functional role of MreB in membrane synthesis has undergone some revision. MreB is now thought to be localized in ‘patches’ and not ‘helices’ as previously thought (Swulius and Jensen, 2012), and in *B. subtilis*, MreB mobility is seen to be determined by the mobility of the peptidoglycan synthesis machinery (Garner et al., 2011). While the molecular interactions between the components of the membrane synthesis machinery have been well documented, how they work spatiotemporally as a system to during membrane growth in *E. coli* remains to be understood.

Here, we examine the spatial growth pattern of single *Escherichia coli* cell membranes using a photobleaching to examine the extent of symmetry in mid-plane growth. We find both membrane growth and nucleoid displacement are asymmetric and correlated. The ‘patch’ distribution and photobleaching recovery MreB-YFP correlates with this growth asymmetry. We attempt to put this in the context of asymmetric growth in other bacteria.

## MATERIALS AND METHODS

### Bacterial strains, plasmids and growth conditions

*E.coli* MG1655 (CGSC 6300) with plasmid based HupA-GFP expression induced with 0.2 % Arabinose (Sisco Research Laboratories, Mumbai, India), as previously described (Wery et al., 2001). MreB was visualized in *E. coli* cells with a YFP-tagged genomic copy using a previously developed strain (Taniguchi et al., 2011) and grown under chloramphenicol selection (Sigma Aldrich, USA).

### Agar micropatterning

Nutrient agar was embossed by altering an approximately 1 mm high layer of molten LB-agar on a PDMS stamp. The stamp has micron-scale ‘H-shaped’ channel pattern designed using CleWin (WieWin, Netherlands). The stamp was made using a two-step soft-lithography approach where a 1 µm layer of gold was sputter-coated followed by spin coating a 2 µm layer of SU8-2 using a spin-coater (Model WS 400B-6NPP/LITE, Laurell Technologies, USA). The photoresist cured with exposure to UV and washed with developer. The exposed gold pattern was washed with acid, leaving a pattern with channels and the trenches (H-pattern) of 2 µm height. A second layer of 20 µm thick SU8-20 was spin-coated and prior to UV-exposure aligned to the lower pattern using the gold. The pattern was then developed. This two-later pattern with 20 µm and 2 µm feature heights was used as a template to make a PDMS stamp by mixing elastomer with curing agent (Sylgard 184, Dow-Corning, USA) in a w/w ratio of 10/1. The stamp was cured by heating it at 60°C in a dry air oven (Raut Scientific, Maharashtra, India) for 2 h. A replica of the PDMS stamp was made by coating an epoxy adhesive mixture (BONDTiTE, Resinova Chemie Ltd., Kanpur, India) with the ratio resin to hardener (w/w) of 8 to 10. Further PDMS stamps were made using the epoxy replica.

### Live cell microscopy

Membrane and nucleoid dynamics were followed by staining a log phase culture of *E. coli* MG1655 cells expressing tagged HupA-GFP with 2 µg/ml of FM4-64 dye (Invitrogen, USA) and incubating for 5 min at 37°C with shaking at 180 rpm. 5 µl of these cells were spread on micropatterned agar pads, incubated at 37°C for 1 hour for cells to recover from the change between liquid and solid medium and mounted on glass coverslip bottomed Petri dishes (60 mm X 15 mm, Corning, NY, USA). Two-channel time-lapse microscopy of these cells was performed on inverted confocal microscope with a plan apochromat 63X 1.40 NA oil immersion objective (LSM 780, Carl Zeiss, Germany) at 1 airy unit using excitation/emission combinations of 633/601 to 759 nm for the FM4-64 channel and 488/493 to 579 nm for HupA-GFP. Images were captured every 11 s for 60 mins while maintaining the chamber at 37°C. Growth of cells imaged in DIC was followed by recording dual channel images in DIC and GFP using the same microscope setup every 2 mins.

### Photobleaching FM4-64 to create membrane landmarks

Alternating cycles of imaging and photobleaching was used follow the landmark of the mid-plane of the cell using a 63X oil immersion lens attached to the confocal microscope described above. Images were acquired every 2 to 4 seconds for 10 to 30 mins and the bleaching to mark the cell midpoint was achieved by selecting an ROI of approximately 0.3 µm wide (along the long-axis of the cell) and increasing the laser power of the 514 nm intensity to 100% for 250 iterations (corresponding to 90 seconds) in order to reduce the fluorophore intensity to 0%. was used for recording images, the magnification was further increased by using optical zoom to varying scales for different fields of interests.

### Analysis of MreB puncta

*E. coli* cells with a genetically modified MreB-YFP were fixed with PFA fixed as described previously (38) and imaged in confocal microscopy with a 63X, 1.4 NA (oil) lens and 514 nm laser for excitation (LSM 780, Carl Zeiss Germany). The data was collected as a Z-stack of 3 µm every 0.1 µm in both YFP and DIC channels and puncta analyzed using a custom ImageJ macro (Code can be downloaded from GitHub https://github.com/Self-OrganizationLab/bactMidcell.git)

### FRAP based estimation of MreB mobility

The MreB-YFP was photo-bleached and the recovery of fluorescence followed. In order to be able to measure kinetics, half the cell of multiple *E. coli* cells were selected and exposed to 50 pulses with 1 s intervals of 100% intensity with a 514 nm laser, to ensure complete bleaching. Images were recorded for next 10 mins at the interval of 1 secs in YFP as well as DIC channels to measure the recovery of fluorescence and cell growth respectively. The fluorescence recovery after photobleaching (FRAP) data was pre-processed in three steps: *(i)* The normalized intensity (*I_norm_*) as a function of time (t) was estimated by using the average intensity in the bleach region (*I_bleach_*) together with the pre-bleach intensity in the same region (*I_prϵ_*) and the intensity of the first post-bleach image that is minimal (*I_min_*) as follows:

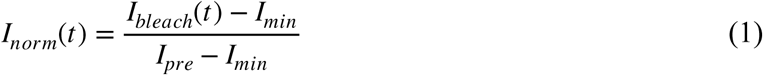

(ii) Correction for image acquisition-based photobleaching of YFP signal was compensated by analysis of a time-series of unbleached cells (Fig. S4A). The intensity profile of such a reference 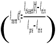 was fit to an exponential decay function with two parameters, a decay rate λ and ‘corrected’ intensity, *I_ref_* as follows:

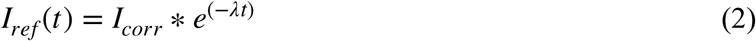

The fit resulted in a decay rate λ of 0.005 1/s. *(iii)* The FRAP intensity 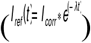 was obtained from dividing the normalized intensity profile obtained from Equation 1 the correction due to exponential decay, as follows:

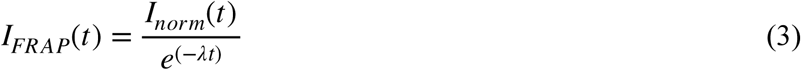

### Image Analysis

The orientation angles of cells on agar pads were quantified by drawing a line was drawn along the length of the cell and measuring the angle between the line and the horizontal axis of the frame using ImageJ 1.50f (Schneider et al., 2012). For tracking movement of the poles and nucleoids, fluorescence time-series images were first stack-registered to compensate for image drift using the ImageJ *StackReg* plugin (P. Thévenaz, U.E. Ruttimann, 1998) and then nucleoids and cell poles were tracked using *ParticleTracker Classic* plugin (Sbalzarini and Koumoutsakos, 2005; Chenouard et al., 2014). The bleach landmark time-series of FM4-64 stained cells were analyzed by constructing a kymograph of either edge of the cell (the bleached region and surrounding membrane from one pole to another) was created using the *MultipleKymograph* plugin and pole growth rates were estimated using ‘*tsp 050706*’ macro. For the MreB puncta analysis, a custom ImageJ macro was developed to extract cell boundaries using the a correlative DIC image and find the geometric mid-plane by finding the midpoint of the spline joining the two poles. The outline was superimposed on corresponding image of MreB-YFP puncta, which were detected using the ‘Analyze Particle’ module of ImageJ (Fig. S6). The macro outputs the ration of sum area of MreB loci to the area of a cellular half in which loci have been detected. Kymographs of DIC image time-series of *E. coli* cells were generated based on interactively selecting an edge of a cell (pole-to-pole) and using the ‘*MultipleKymograph*’ plugin with the ‘*tsp050706*’ macro in ImageJ, as before.

### Data analysis

All the data was plotted and analyzed using MATLAB R2010b (Mathworks Inc., USA) and GraphPad Prism 5 (GraphPad Software, CA, USA) respectively. All the figures were prepared using Inkscape 0.91 (open source software GPL, https://inkscape.org).

## RESULTS

### Aligned growth of *E. coli* cells in patterned agar-pad trenches

Cell alignment in culture was considered an essential step in successful analysis of the growth dynamics of single cells. In order to achieve this, we have grown them on a solid medium in channels. The solid medium prevents diffusive motion seen potentially in microfluidics systems of single-cell analysis. This could potentially confound the growth assessment. The growth of cells in channels was aimed to avoid more than one cell growing in contact with another, by playing with the sensing density. To this end, micro pattern based masks were constructed using soft-lithography and epoxy replicas with a length: 1 mm, depth: 2 µm and width: 1 to 20 µm. Onto these masks, nutrient agar was deposited in a thin layer of approximately 5 mm (Fig. 1A). The agar pad obtained by this manner had indentations that can be visualized in a dissection microscope (Fig. 1B). A time-series of cells grown in the region with 1 µm wide channels (Fig. 1C, Video S1) was analyzed for the degree of alignment of cells, using the long-axis of the cell and an alignment of 15 to 24 degrees appears to have been achieved, based on three representative time points (Fig. 1D). The alignment angles of cells in the trenches are comparable and the used further to analyze cell elongation of the pairs of poles.

**FIG 1.**
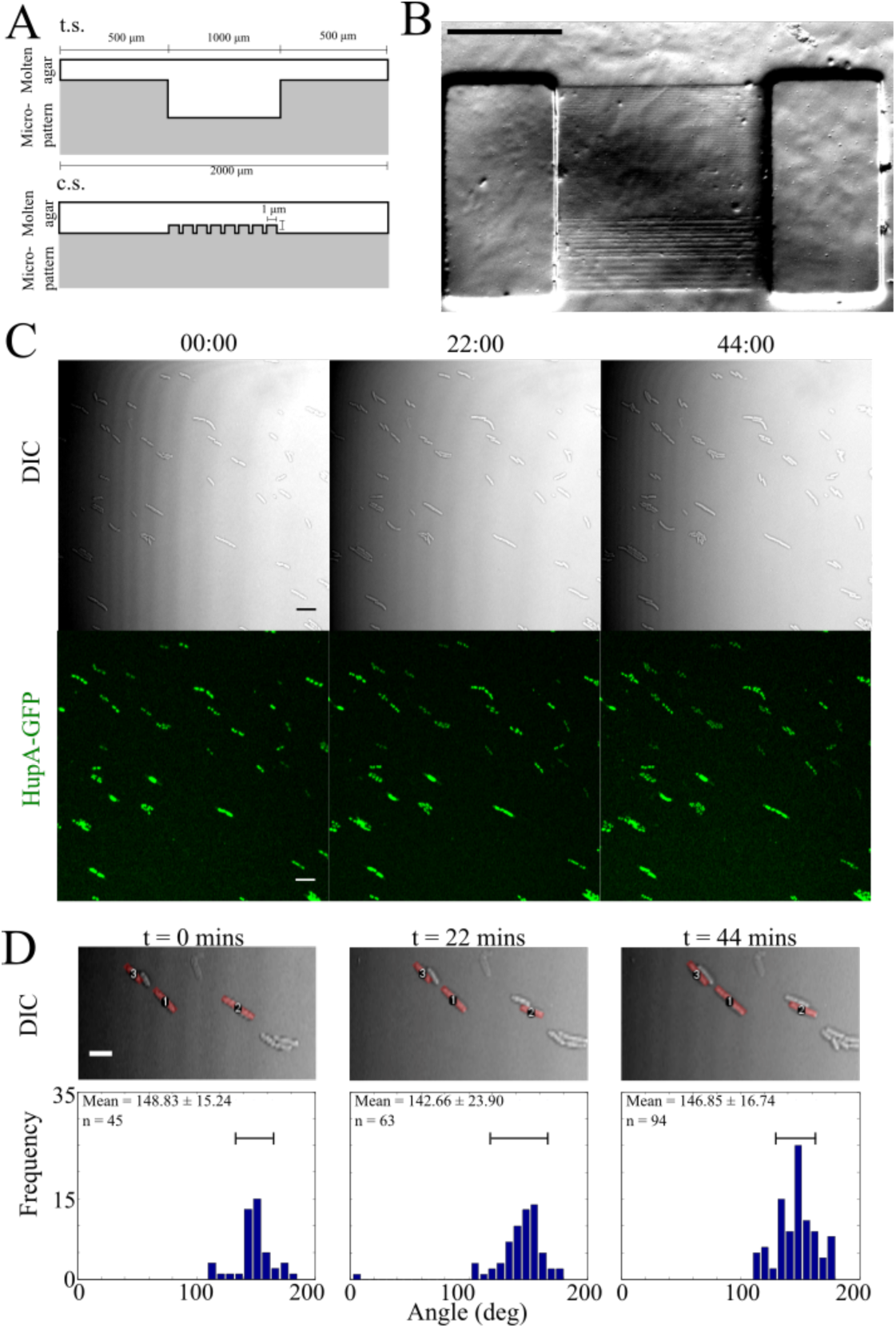
Spatially confined growth of E. coli cells. (A) The micro-patterned agar pad is polymerised on a PDMS stamp with 1 µm high and wide channels (t.s.: transverse section; c.s.: cross section). (B) A bright-field image of the final micro-patterned agar pad (scale bar: 500 µm). (C) Representative images of a time-series of *E. coli* MG1655 cells in DIC and fluorescence to follow HupA-GFP tagged nucleoid. Time stamp: hh:mm. Scale bar: 5 µm. (D) Representative DIC images of cells on the agar pad overlaid with the long axis (red line), used to estimate the orientation angle taken at multiple time points (t). Scale bar: 5 µm. The frequency histogram of the angle of cells with the positive x-axis of the image. The bar marks the mean +/− s.d.

### Asymmetric mid-plane membrane addition during *E. coli* cell growth

*E. coli* cell wall synthesis during cell growth has been recognized to involve active growth sites spread along its length away from the cell poles (Scheffers and Pinho, 2005). FM4-64 is used to label the membrane and the mid-plane region is marked by periodic photobleaching of the FM4-64 dye (Fig. 2A,B, Video S2). The differences in the growth of two cellular halves is visible in kymographs of cellular growth constructed from the contour along the fluorescently labelled cell membrane, along the cell length (Fig. 2C). The apparent drift of the reference point (red line) away from the fast growing end of the cell is interpreted as a gradual deviation of bleached ROI from mid- plane, due to the growth bias, resulting in a new geometric cell centre in the course of growth. By choosing cells free of potential pushing forces from neighbors, we believe we have measured intrinsic growth asymmetry and not translation of cells (Fig. S2, Video S2). The instantaneous growth rates, obtained from the slope of the bleach marks in the kymographs between the two successive rounds of bleach pulses, demonstrate quantitative differences in membrane addition between ‘slow’ and ‘fast’ growing cell halves or ‘poles’ (Fig. 2D). The difference between pairs of instantaneous growth rates was found to be 0.0628 µm/ min (n= 18 cells) (Fig. 2E). However, such asymmetric cell growth would be inconsistent with the high degree of precision reported during equational division in *E. coli* cells. To test whether this asymmetric growth is compensated for by the genome positioning, we proceeded to follow nucleoid segregation dynamics.

**FIG 2.**
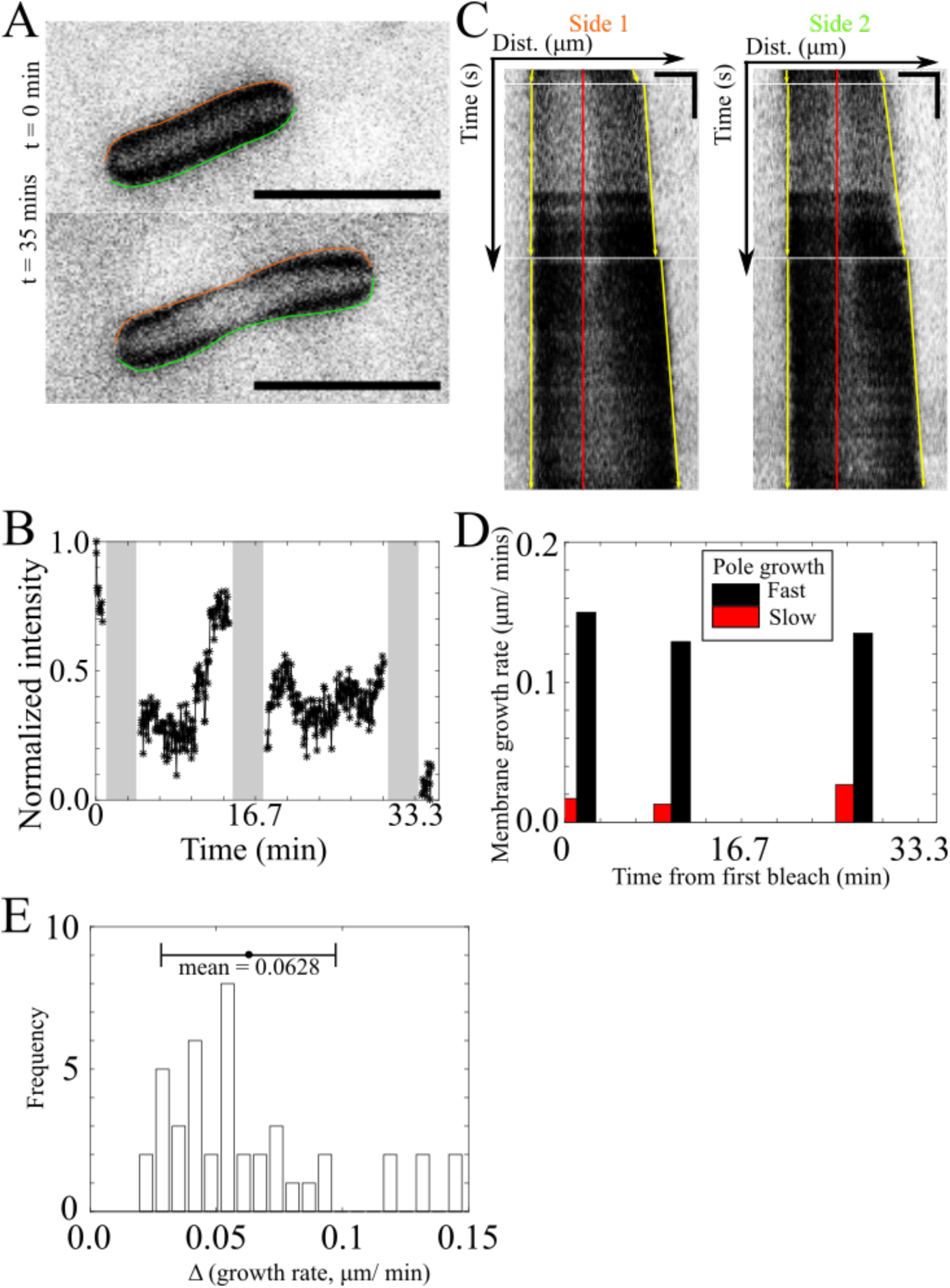
Asymmetric growth of the membrane. (A) FM4-64 labelled *E. coli* cells at time 0 and 35 min (scale bar: 5 µm) with the contour of the cell membrane marked (side 1: red side 2: green). (B) The normalized fluorescence intensity of the mid-cell region is plotted as a function of time. Grey boxes indicate periods of photo-bleaching. (C) Two representative kymographs (sides 1 and 2) generated from the same cell in (A) (scale bars: 2 µm, 150 s). Horizontal white lines: bleaching periods, yellow line: edge of bleach-mark. (D) The rates of bleach mark expansion between bleach periods for one cell and (E) the frequency distribution of the absolute difference between expansion rates from multiple cells (n = 18).

### Nucleoid displacement correlates with a membrane growth

In order to examine how growth asymmetry could be compensated for by genomic DNA positioning, we simultaneously tracked the displacement of the cell pole based on the FM4-64 membrane label (Fishov and Woldringh, 1999), and plasmid expressed HupA-GFP to label the nucleoid (Wery et al., 2001). We examined if the displacement of the two daughter nucleoids and their respective poles could be correlated (Fig. 3A). Mean nucleoid segregation rates varied between 0 and 0.08 µm/min while cell pole displacement rates varied between 0.02 to 0.09 µm/min (n = 13 cells) with a strong linear correlation between the two (Fig. 4B). The mean difference between the pairs of cell poles and daughter nucleoid segregation rates also appeared to be linearly correlated (Fig. 4C). These observations are consistent with the equational division, despite asymmetric membrane growth. We proceeded to examine whether a bacterial cytoskeletal protein MreB, that mediates cell membrane growth, might also be correlated to the observed growth bias.

**FIG 3.**
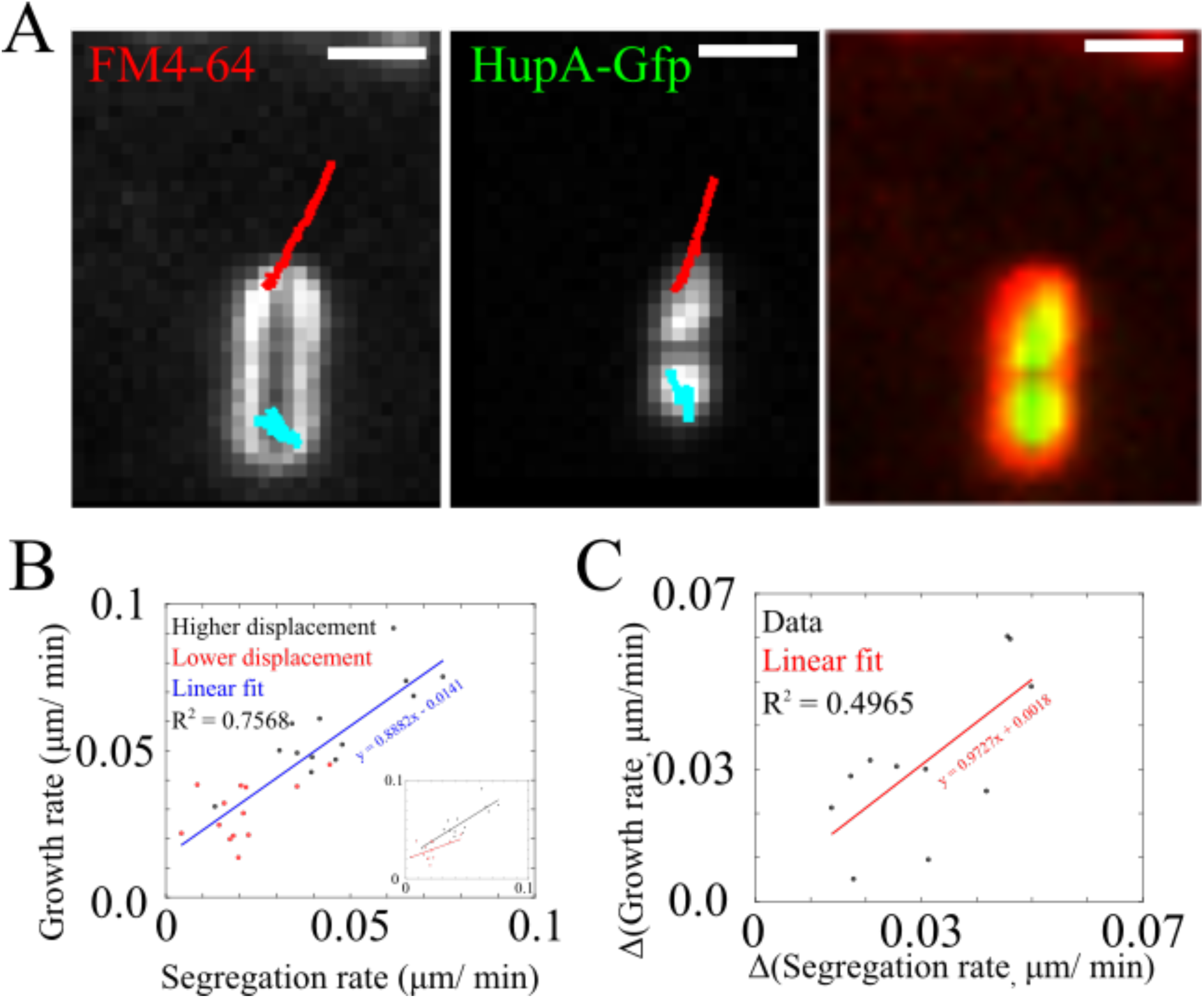
Correlated asymmetry in nucleoid segregation and pole displacement. (A) *E. coli* cell poles and nucleoids were tracked over time (red, cyan) based on FM4-64 staining and HupA-GFP expression respectively as seen in the first frame of the time series from the two channels and the merge image (red: FM4-64, green: HupA-GFP.) Scale bar: 2µm. (B) The net displacement rate of the fast (black) and slow-growing poles (red) is plotted as a function of the segregation rate of the respective nucleoids for multiple cells and (C) the difference between the growth rates of poles and the segregation of daughter nucleoids.

**FIG 4.**
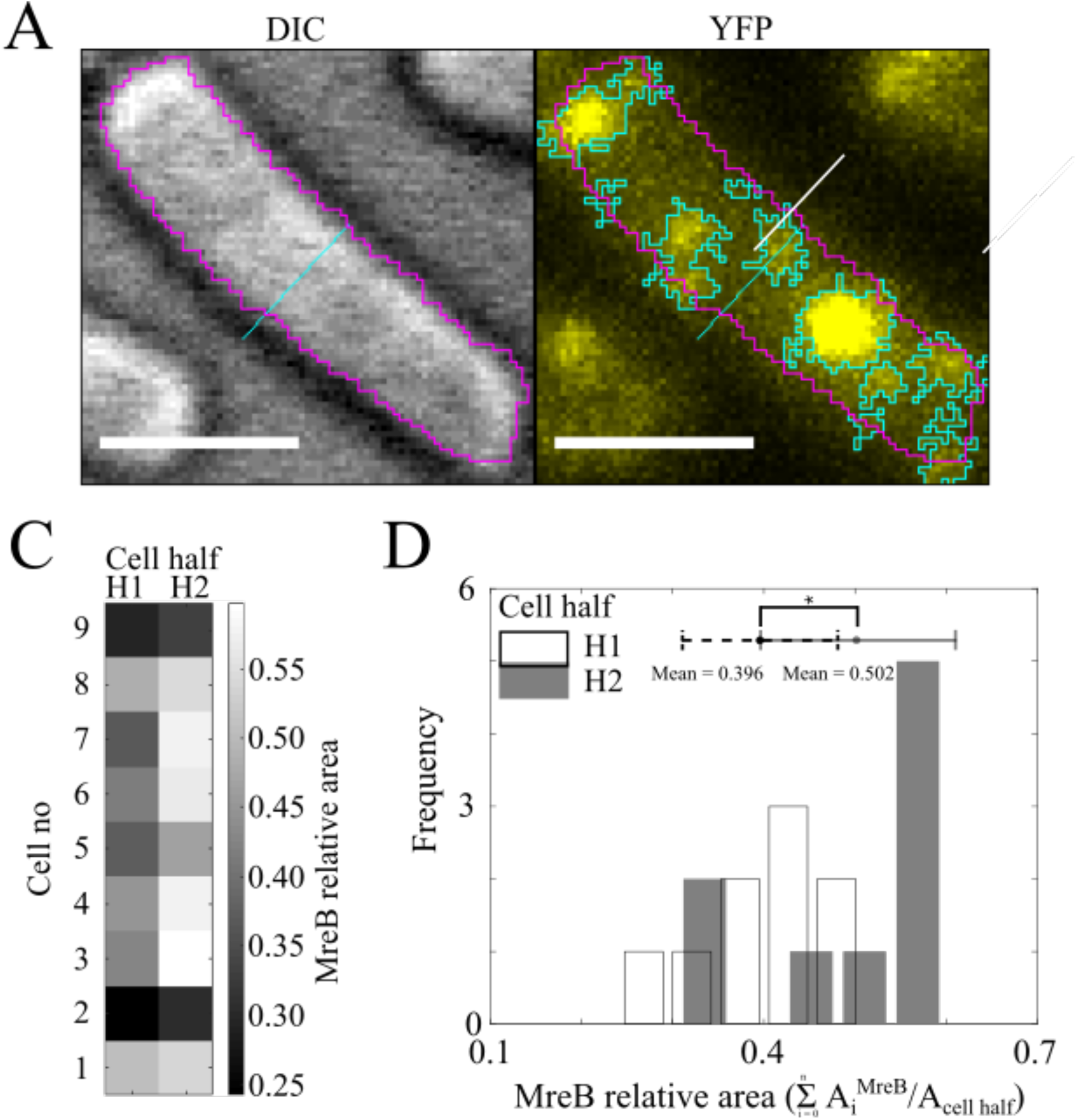
Asymmetric distribution of MreB foci. (A) The cell outline (pink) and mid-plane (cyan) were identified from DIC images of *E. coli* and the coordinates applied to the fluorescence channel of the same cell where MreB-YFP puncta were detected (cyan). This was used to obtain the area occupied by MreB in each cell half. Scale bar: 2 µm. (C) The relative area occupied by MreB (colorbar) in each cell half was classified as either sparse (H1) or dense (H2) for 9 cells that were analyzed. (D) The mean area fraction for pairs of cell halves (H1 vs. H2) is significantly different (t-test with n = 9, df = 16, p = 0.0243). The mean area fraction in the sparse half (H1) is 0.396 and dense half (H2) is 0.502.

### Patch distribution and FRAP recovery kinetics of MreB

The role of MreB in the regulation of cell wall architecture as well as in the maintenance of cell shape (Carballido-López, 2006) suggests that studying the dynamics localization of this polymer could correlate with the growth asymmetry. Fixed *E. coli::mreB-yfp* cells (Taniguchi et al., 2011) were imaged in a confocal microscope and the punctuate localization of MreB protein quantified (Fig. 4A) using ImageJ (Fig. S3). The area occupied by MreB-YFP puncta in the two cell halves appears asymmetric i.e. the total area occupied by MreB puncta in H1 is greater than area in H2. Here Hi (i=1 or 2) represents the geometric half of the cell. We adopt a convention where i=1 is used to indicate the cell-half with the larger area. The distributions of MreB puncta in either halves of *E. coli* cell appear to be significantly different based on a T-test (n = 9, df = 16, p = 0.0243) (Fig. 4D). In order to further test this observation the mobility of MreB in a cell half was also examined.

The fluorescence recovery after photobleaching (FRAP) of MreB-YFP in *E. coli* cell halves was then correlated to DIC image based pole-displacement as a measure of cell growth (Fig. 5A). Representative kymographs suggest the faster growing cell half also demonstrates faster recovery dynamics of MreB (Fig. 5B). In order to quantify the dynamics, FRAP dynamics were analyzed in three steps: (a) normalization by the pre-bleach (maximum) and bleached (minimum) intensities (Equation 1), (b) estimating the degree of imaging based photobleaching in a reference cell that is not bleached (Equation 2) and (c) correcting for imaging based photobleaching. (Equation 3). These corrected FRAP intensities, *I_FRAP_* (Equation 3), were fit to single exponential function to calculate mobile fraction (F_mob_) and half-recovery time (τ) as follows:

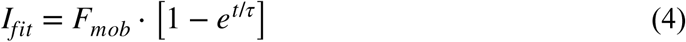

**FIG 5.**
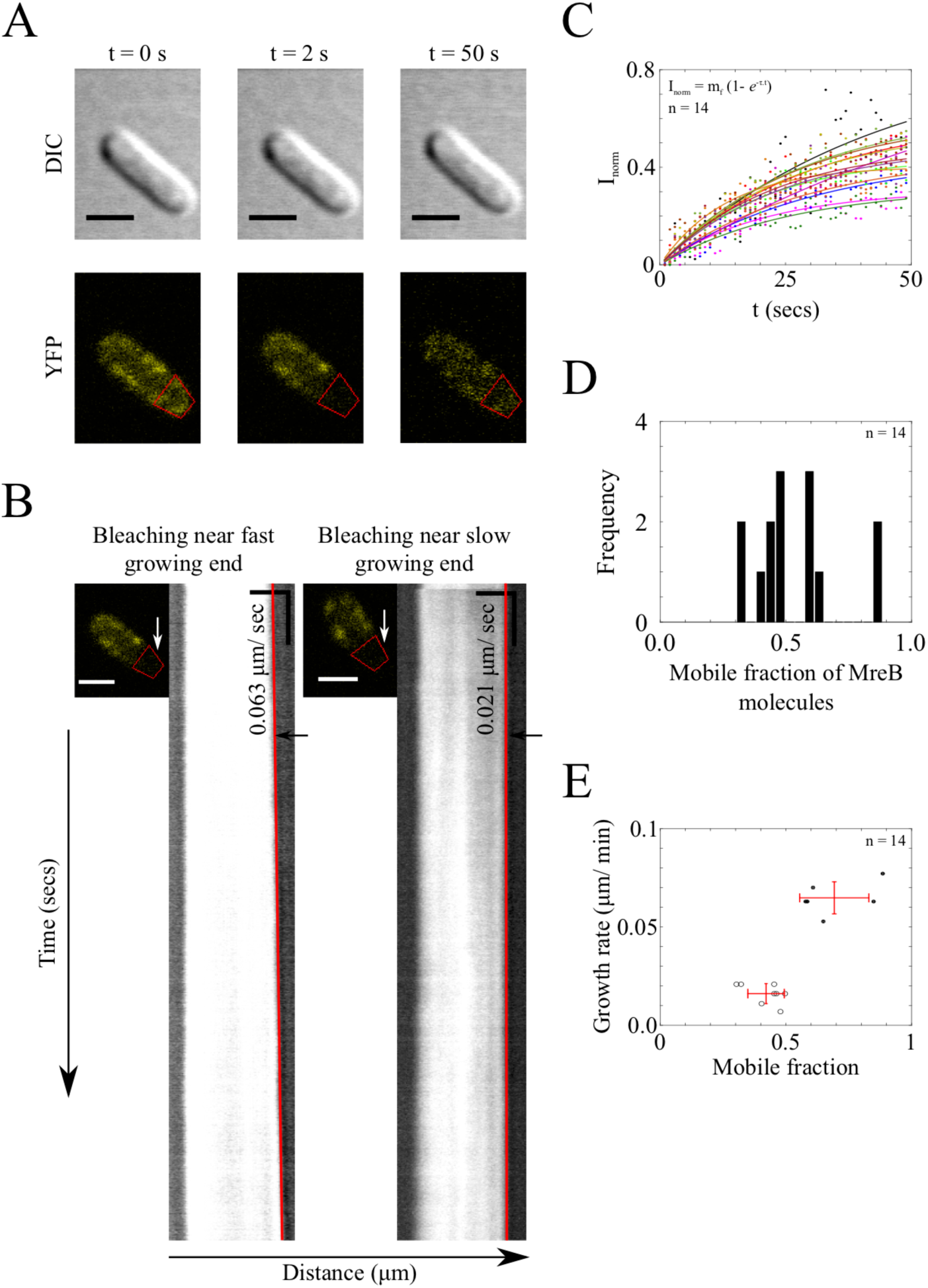
MreB FRAP mobility in cell half correlated to membrane growth. (A) *E. coli* cell with endogenous MreB YFP (*lower left panel*) was bleached near one of the poles (*lower middle panel*) and observed for fluorescence recovery (*lower right panel*) in YFP and in DIC (*upper panel*) channels. Scale bar-2 µm. (B) Kymographs from DIC time-series of *E. coli* are used to estimate estimate the growth rate of the cell halves based on the slop of the each cell pole (solid red line). Scale bars-2 µm (dist.) and 50 s (time). (C) Normalized YFP intensities were fit to the single exponential function to determine mobile fraction and half recovery time of MreB molecules in the bleached region (n = 14 cells, represented by different colors). (D) The frequency distribution of MreB mobile fractions from the regions near pole and (E) the plot of growth rates of the bleached cell-half plotted against the respective the mobile fraction of MreB. The growth rates (+/−s.d.) and the mobile fractions (+/−s.d.) were subjected to an unpaired t test (n = 14, df = 12, growth rate p < 0.0001, Mobile fraction p = 0.0004).

The fits to the MreB-YFP half-cell FRAP data (n = 14 cells) demonstrate a range of recovery kinetics (Fig. 5C). None of the curves reach the maximal value of 1, suggesting a sizable fraction of the proteins are immobile. This is consistent with the polymerized nature of MreB. The mobile fraction obtained from the fits (Equation 4) appears to suggest multiple clustered values (Fig. 6D). To our surprise, when the mobile fractions were correlated to the growth rates of the respective poles, falling into two distinct groups. Cell halves with F_mob_ ≥ 0.5 correspond to faster growing cell poles while F_mob_ = 0.3 to 0.5 were associated to slow growing cell poles (Fig. 5E, Table S1). The average of half recovery time (t_1/2_) of MreB-YFP was found to be 20.1 secs (see Fig. S5A) and it correlated linearly with increasing mobile fraction (Fig. S5B). The t_1/2_ was also correlated to the pole growth rate (Fig. S5C), similar to the trend in F_mob_. Our observations, thus suggest a potential role for MreB, the actin-like cytoskeletal protein in asymmetric cell growth.

## DISCUSSION

Here, we have demonstrated that membrane addition at the mid-plane of *E. coli* appears to result in asymmetric elongation of the geometric centre of a newborn cell. At the same time, the linearly correlated asymmetry in nucleoid displacement suggests how such an asymmetry still results in geometrically equal cell halves with equidistant positioning of the nucleoids from the cell poles. Additionally we find the distribution and dynamics of MreB appear to correspond to faster growth, suggesting a potential link between the dynamics of this cytoskeletal element that drives membrane addition and cell growth asymmetry.

Bacterial growth at just one pole has previously been observed for various prokaryotes such as *Mycobacteria* that lack MreB protein and grow at the one pole before dividing (Aldridge et al., 2011; Joyce et al., 2012). On the other hand, asymmetric growth is observed in *Caulobacter crescentus* culminates in asymmetric division and differentiation (Skerker and Laub, 2004). Asymmetry in growth, however, has been seen as a mechanism exploited by bacteria in order to restrict damage to only a few cells in the population, also referred to as ‘cellular aging’, eventually culminating in the death of those cells that accumulate large amounts of damaged material (Stewart et al., 2005). Although an array of processes ensure the symmetric placement of division site in *E. coli* (Raskin and De Boer, 1999; Varma et al., 2008), the duplication of *E. coli* cell has been observed more recently to result in daughter cells, whose molecular compositions are distinctly different due to accumulation of protein aggregates preferentially in the old pole (Lloyd-Price et al., 2012). Our study complements such observations with evidence for spatial asymmetry arising from uneven growth along the mid-plane. It would indeed be interesting to see if the aggregates correlate with the membrane growth asymmetry.

Very early models of cell size growth of *E. coli* had assumed surface extension as a symmetric growth process, with active growth at the poles (Doanachie, 1977). However, more recent studies on membrane growth have demonstrated the mid-plane addition of membrane (Rojas et al., 2018; Ursell et al., 2012). Our observations of growth in the cell mid-plane however suggest the rate of membrane growth in either direction is not identical. Our analysis included cells proliferating with different growth rates. We report the average growth rate due to membrane addition of the fast growing cell half is 0.05-0.08 µm/min, while the other half grows with an average speed of 0.01- 0.04 µm/ min. While these differences are extremely small, they appear to reproducible.

The use of a bleach-mark in the FM4-64 staining of the membrane, results in recovery of the signal, due to new membrane addition and mobility of the membrane. As a result, we repeatedly bleached the region. The asymmetric growth of the cell results in the formation of newer mid-plane which was reflected as a clear deflection of the bleached region in a kymograph generated as a temporal interpretation of LOI drawn on the lateral side of the cell along the length. For few cells we also observed decrease in the fluorescence intensity at the growing arm over time (data not shown). Incorporation of more lipids in the growing end dilutes the dye concentration in that region because of which the intensity of FM4- 64 on either side of the bleach mark becomes unequal. In addition to this, kymograph showed two different slopes for two poles, indicating different growth rates. Our experiment not only helped us to confirm our observations but also provided with a novel technique to create a spatial marker on bacterial surface.

The asymmetric distribution of the growth along the cell length imparts an obvious asymmetry in the segregation dynamics of sub- cellular components. The measurements of nucleoid displacements in a pair conceded with our hypothesis. Though the method used to estimate the nucleoid segregation speed was not entirely objective, the tally of average nucleoid speed we obtained (0.033 µm/ min) from the analysis with previously reported value (Nielsen et al., 2006), established the reliability of the results. Our studies of difference in the displacements of sister nucleoids with respect to the growth asymmetry revealed a linear relationship between the distance travelled by a nucleoid and the growth of its nearest pole, implying that one of the sister chromosomes is compelled by higher growth at its nearest end to cover more distance as compared to its pair before it is located to the sub- cellular compartment. Importantly, the difference between the nucleoid segregation rates, when plotted against the difference between the corresponding pole growth rates, form a line with a slope approximately equal to 1. We believe that the extra distance added in the nucleoid movement is equivalent to the additional growth at its corresponding end, permitting the cell to eventually divide in the geometric centre.

The rod shaped microorganisms like *Mycobacterium sp.* and *Corynebacterium sp.* lack the MreB gene. In these organisms the growth is guided by the active pole (Aldridge et al., 2011; Daniel and Errington, 2003). In the round cells, it has been shown that FtsZ anchors PBP 2 to the periphery near the poles and initiates the cell wall synthesis (Pinho and Errington, 2003). We report for the first time the dynamics of MreB molecules and correlate it with asymmetric growth. To analyse the area occupied by MreB in each cellular half, we developed an ImageJ macro. The macro uses DIC images to extract the outline of the cell. The outline is used to find the cell contour as well as the mid- plane of the cell. The cell is thus divided into two halves and subsequently the two ROIs representing two cellular halves are overlaid on the corresponding fluorescence image of MreB- YFP. The total area covered by MreB puncta is then estimated separately for each half and represented as the sum area of MreB foci normalized by the area of respective cellular half (see Fig. S6). Though we developed a macro to assess MreB localization, it could be more widely used.

The MreB distribution is uneven across the cell length. Our estimate was further corroborated by FRAP studies in live *E. coli* cells. We observed a threshold value of 0.5 in mobile fraction. The concentration of MreB molecules was found to be more near the fast growing end. At the same time, its mobile fraction is less than 0.5 near the pole with slow growth rate. MreB is an important molecule in *E. coli* growth, as it provides a scaffold for the growth as well as controls the cell wall construction through the orchestration of cell wall synthesizing machinery (Carballido-López, 2006; Errington, 2015). Heterogeneous distribution of MreB molecules across the longitudinal axis of the cell explains the asymmetric growth of an organism.

The growth asymmetry in organisms has been associated with aging. In rod shaped single celled organisms, septum is converted into the new pole, thereby, making the existing pole older one. Previous examinations of the rod cells have revealed that the defects are majorly compartmentalized towards the old pole (Stewart et al., 2005). In *Mycobacteria* old poles are considered as leading pole at which the maximum growth takes place. The pattern is repeated through the generations (Aldridge et al., 2011). Similar studies in *E. coli* confirm the inheritability of growth asymmetry through consecutive generations. Though our studies were limited to two generations due to consequent dilution of FM4- 64, we found that the inheritability pattern in *E. coli* differs from *Mycobacteria*. Unlike Mycobacterium, new born *E. coli* daughters do not exhibit the growth bias towards old pole. The daughter with fast growing end from the mother cell, initiates its growth right after septum formation and grow towards the old pole. The slow growing end of the mother cell turns out to be the slow growing pole in the next generation. In such daughters, new pole develops into fast growing end. However, we observed that the later cell always lags in the growth, generating difference in the doubling times of the two sisters (data not shown). Though the phenomenon has been observed in *Mycobacterial cells* (14), it did not give rise to phenotypic heterogeneity in *E. coli* populations. We reasoned that Mycobacterial cell cycle is regulated by ‘timer’ mechanism (Aldridge et al., 2011), on the other hand BCD cell cycle of *E. coli* follows ‘adder’ mechanism (Taheri-Araghi et al., 2014). Hence, the growth lag in one of the sisters does not cause premature division.

In conclusion, we investigated the growth in single cell of an *E. coli*, using three different techniques. Our work suggests the presence of a slight asymmetry in *E. coli* membrane addition correlated with nuclear displacement, which is in contrast with classical model for *E. coli* cell growth.

## ACKNOWLEDGEMENT

The pBAD24-HupA-GFP plasmid was a kind gift from Dr. Josette Rouviere-Yaniv. We would like to thank Dr. Sunish Radhakrishnan for critical feedback.

The Authors declare that there is no conflict of interest.

MSG and CAA designed the study. MSG performed the experiments with bacteria, data analysis and made the figures. CAA performed the experiments for micro-patterning and designed the analysis. CAA wrote the paper.

## SUPPLEMENTAL MATERIAL

**FIG S1.**
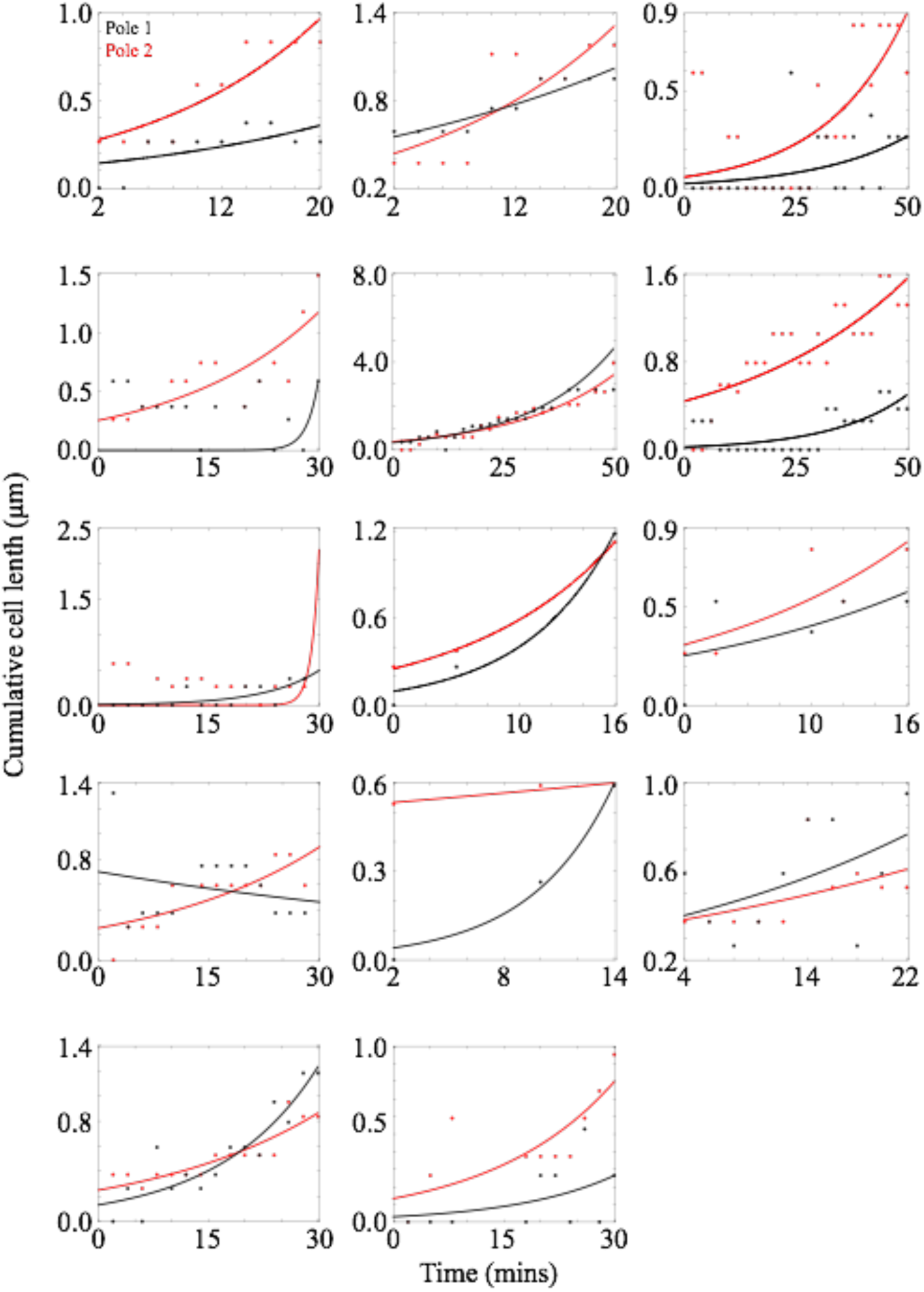
Growth rates of the cell poles were quantified by fitting exponential function to the cumulative displacements exhibited by the poles. Growth near polar region at each time point was estimated by Euclidian distance formula with respect to the initial time point. The cumulative pole growths (black and red filled circles) thus obtained for 14 cells were then fit to exponential fits (black and red solid lines) to estimate the growth rates of an individual end of the cell.

**FIG S2.**
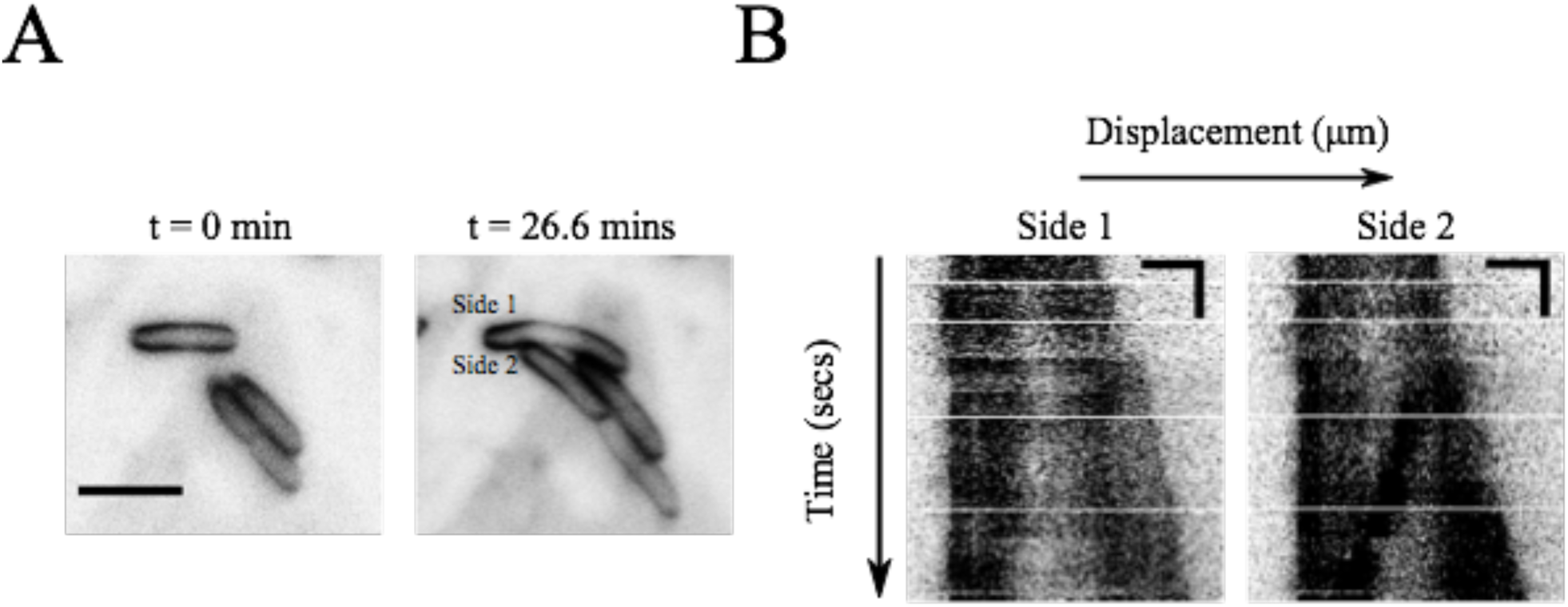
Growth of cells with close (A) Representative images of *E. coli* cell stained with FM4-64 (inverted) have been shown for t = 0 mins and t = 35 mins. (B) Kymographs (inverted) generated for both the sides of the cell around its major axis have been depicted. (Horizontal scale bar- 2 µm; vertical scale bar- 100 secs).

**FIG S3.**
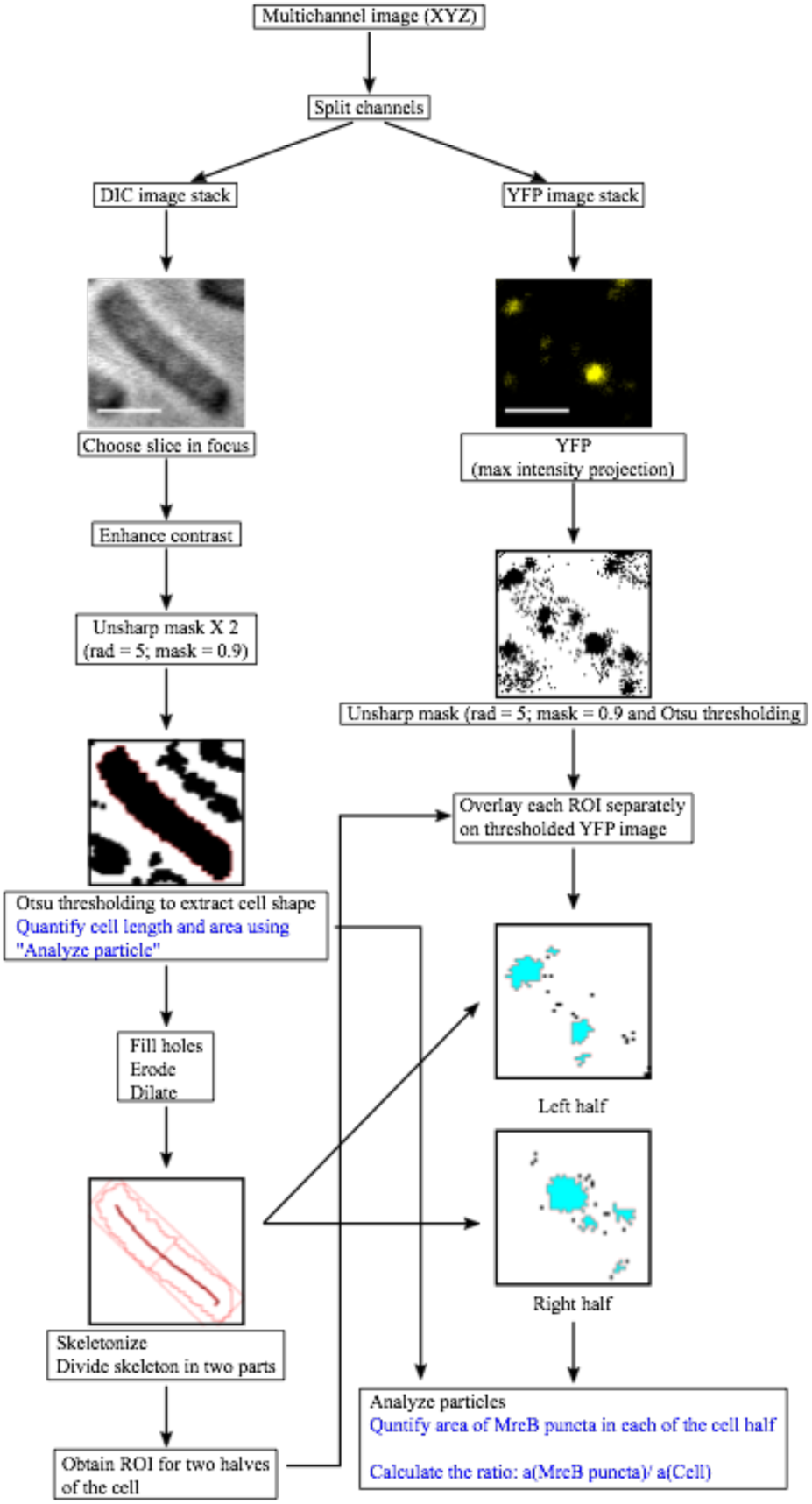
Spatial analysis of MreB puncta distribution in the cell. Fixed cells of *E. coli* were imaged for MreB puncta localization in DIC and YFP channels. DIC image was processed to extract the parameters like cell shape, area, length and geometric center. Using cell center two ROIs representing each half of the cell were obtained. Each ROI was separately overlaid on YFP image to detect the MreB puncta in defined area. Area of all detected MreB loci was summed up and its ratio was taken over the total cell area.

**FIG S4.**
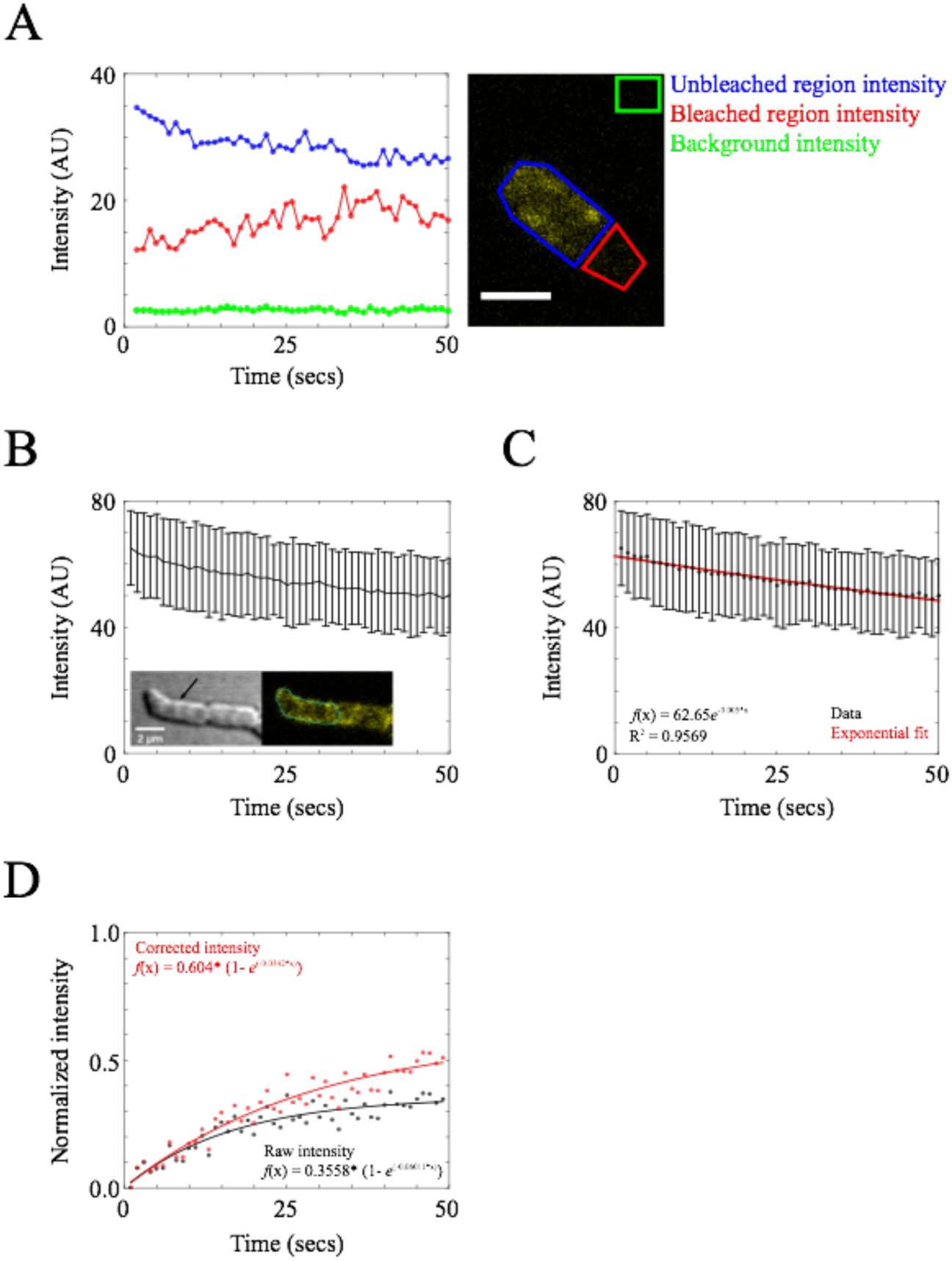
Intensity values for post- bleaching period were corrected for YFP photo- bleaching. (A) Intensities from bleached region (red), unbleached region (blue) of the cell and from background (black) have been plotted for first 50 secs of the experiment (B) Intensity values from reference region were plotted as function of a time. Inset shows the DIC as well as YFP snapshots of the representative reference cell (indicated by arrowhead) at t = 0 secs. The reference region has been marked with cyan boundaries. (C) The average intensities obtained from 11 reference cells were fit to an exponential decay equation to calculate half decay rate (0.005 s^−1^) (D) Normalized intensities of bleached region before correction (black) and after correction (red) were plotted for first 50 secs and fit to single exponential function.

**FIG. S5.**
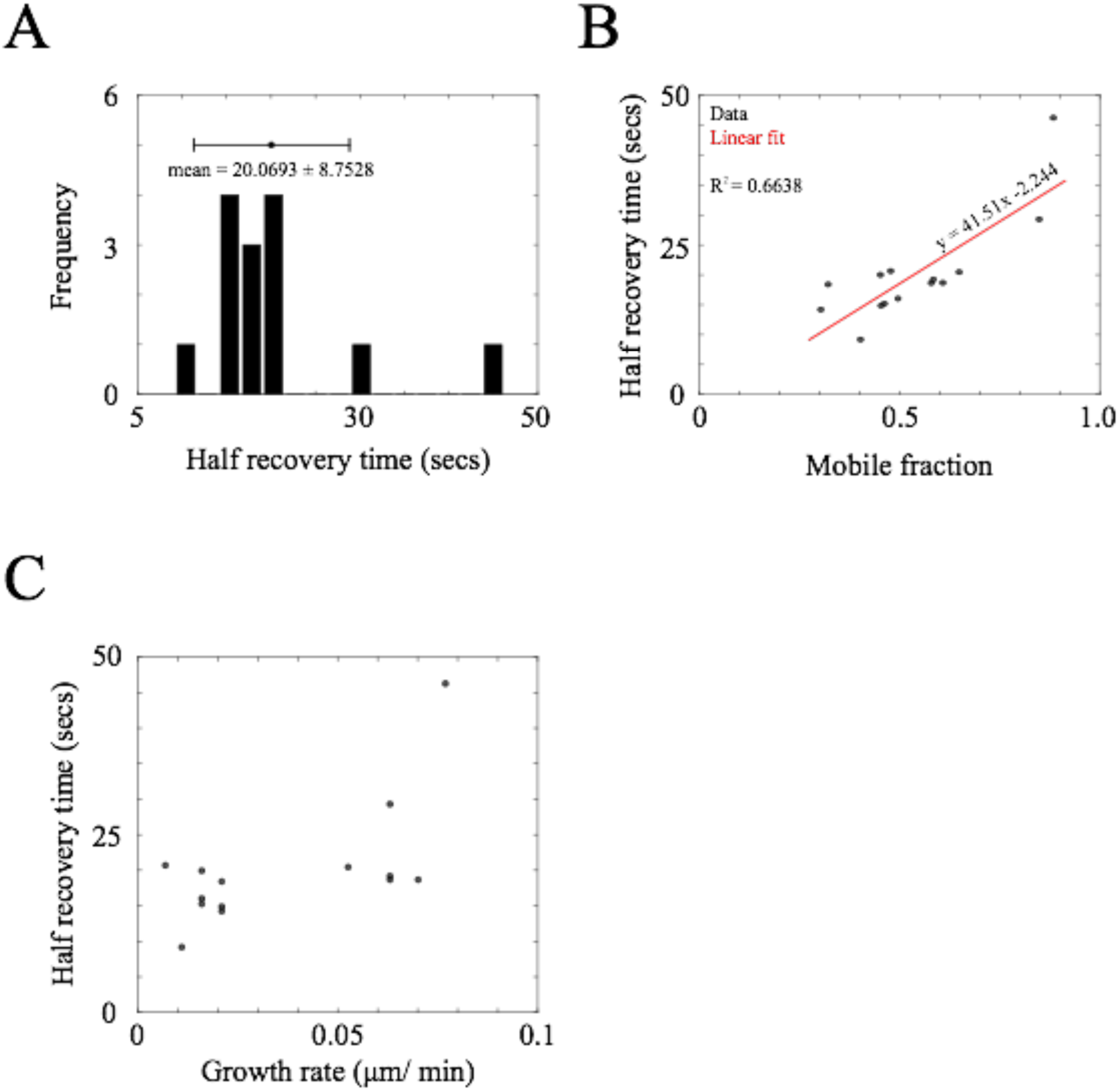
Half recovery time of MreB YFP molecules is independent of their mobile fraction. (A) The distribution of half recovery time of MreB YFP molecules obtained after bleaching *E. coli* cell (n = 14) has been depicted (B) The half recovery times has been plotted as the function of corresponding mobile fractions of MreB-YFP molecules in the bleached cellular end (n = 14) and are fit to the linear function (solid red line) (C) The half recovery times are plotted against the growth rate of their nearest pole.

**Table S1.** End-growth rates of bleached and unbleached ends of growing *E. coli* cell and correlation with the mobile fraction and half recovery time at the bleached end. The highlighted cells show higher mobile fraction near fast growing end of the cell.

| Growth rate of the bleached end ($\mu\text{m}/\text{min}$) | Growth rate of the unbleached end ($\mu\text{m}/\text{min}$) | Mobile fraction of MreB YFP | Half recovery time (secs) |
| --- | --- | --- | --- |
| 0.063 | 0.005 | 0.8490 | 29.27 |
| 0.063 | 0.007 | 0.5846 | 19.23 |
| 0.07 | 0.028 | 0.6082 | 18.67 |
| 0.063 | 0.007 | 0.5794 | 18.64 |
| 0.077 | 0.014 | 0.8844 | 46.21 |
| 0.0527 | 0.095 | 0.649 | 20.45 |
| 0.021 | 0.056 | 0.3034 | 14.2 |
| 0.007 | 0.049 | 0.4779 | 20.64 |
| 0.021 | 0.059 | 0.4539 | 14.85 |
| 0.016 | 0.037 | 0.4962 | 16.03 |
| 0.021 | 0.047 | 0.3216 | 18.39 |
| 0.011 | 0.058 | 0.4025 | 9.2 |
| 0.016 | 0.053 | 0.4523 | 20.0 |
| 0.016 | 0.063 | 0.4623 | 15.19 |

**Video S1.**
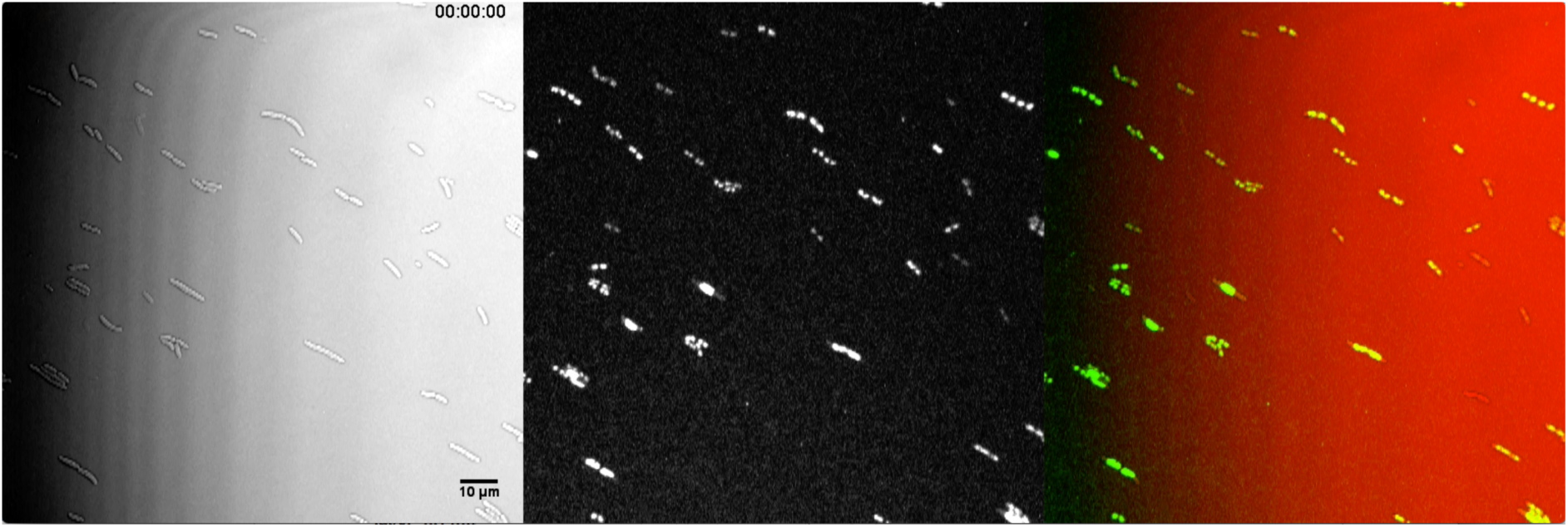
Growth of *E. coli* MG1655 cells transformed with tagged extra-chromosomal copy of HupA on micro-patterned agar pad. Time lapse images of live *E. coli* MG1655 cells growing on LB agar were captured at the time interval of 2 mins in DIC (*left panel*) and GFP (*middle panel*) channels using confocal laser microscope (LSM 780; Carl Zeiss). Scale bar- 10µm. Time lable- hr: min: sec.

**Video S2.**
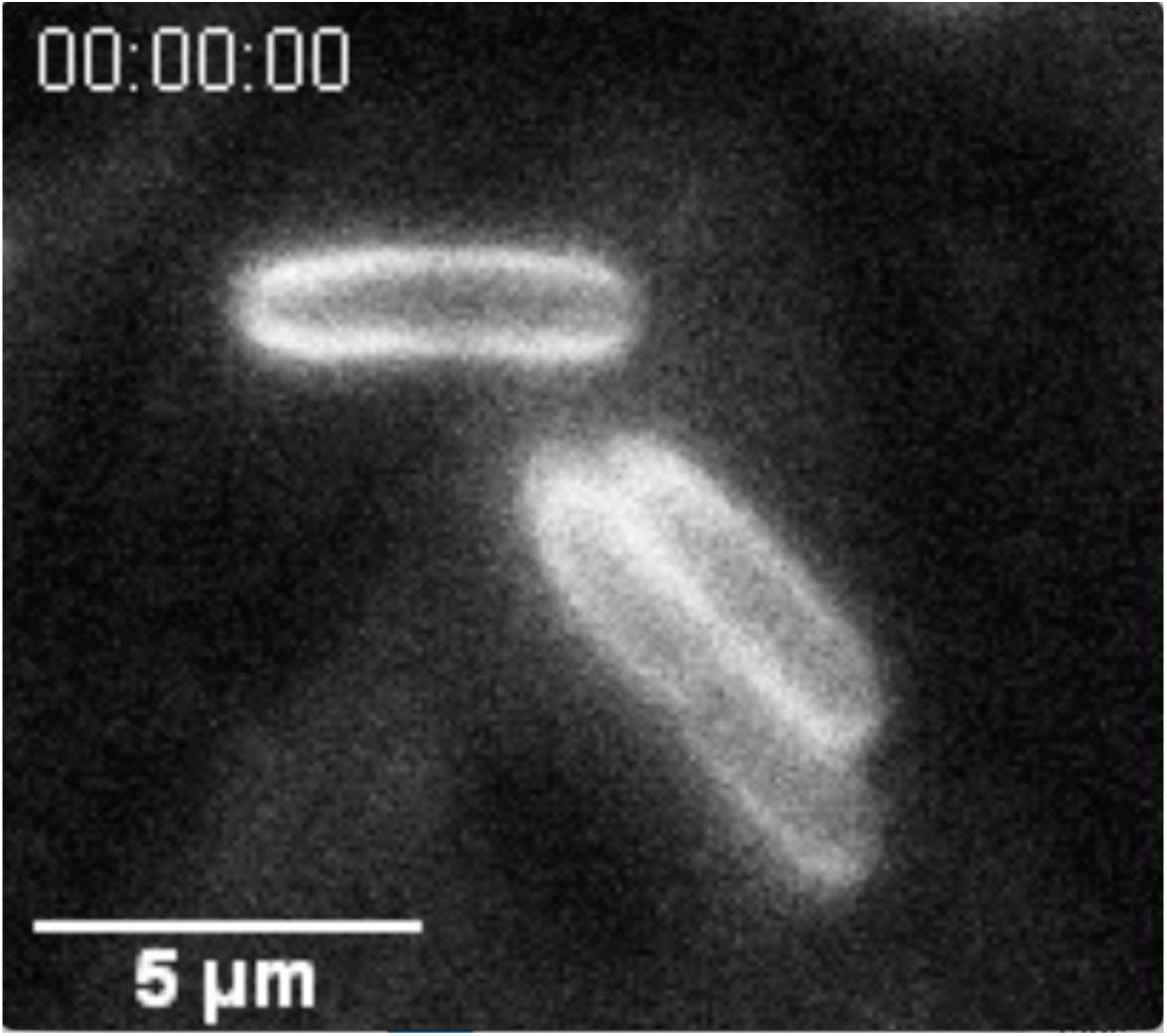
Growth of *E. coli* cell stained with FM4- 64 bleached at its mid- plane. An *E. coli* cell membrane stained with FM4- 64 is bleached at its mid-plane and imaged at the regular interval of 4 secs. White arrows indicate the bleached regions. Scale bar- 5 µm. Time label- hr: min: sec.

## References

Aldridge, B.B., M. Fernandez-suarez, D. Heller, V. Ambravaneswaran, D. Irimia, M. Toner, and S.M. Fortune. 2011. Asymmetry and Aging of Mycobacterial Cells Lead to Variable Growth and Antibiotic Susceptibility. Science. 335:100–104.

Beall, B., and J. Lutkenhaus. 1991. FtsZ in Bacillus subtilis is required for vegetative septation and for asymmetric septation during sporulation. Genes Dev. 5:447–455. doi: 10.1101/gad.5.3.447.

Ben-Yehuda, S., and R. Losick. 2002. Asymmetric Cell Division in B. subtilis Involves a Spiral-like Intermediate of the Cytokinetic Protein FtsZ. Cell. 109:257–266.

Brown, P.J.B., M.A. De Pedro, T. Kysela, C. Van Der Henst, J. Kim, X. De Bolle, C. Fuqua, and Y. V Brun. 2012. Polar growth in the Alphaproteobacterial order Rhizobiales. Proc. Natl. Acad. Sci. 109:3190–3190. doi:10.1073/pnas.1200309109.

Carballido-López, R. 2006. The bacterial actin-like cytoskeleton. Microbiol. Mol. Biol. Rev. 70:888–909. doi:10.1128/MMBR.00014-06.

Chenouard, N., I. Smal, F. de Chaumont, M. Maška, I.F. Sbalzarini, Y. Gong, J. Cardinale, C. Carthel, S. Coraluppi, M. Winter, A.R. Cohen, W.J. Godinez, K. Rohr, Y. Kalaidzidis, L. Liang, J. Duncan, H. Shen, Y. Xu, K.E.G. Magnusson, J. Jaldén, H.M. Blau, P. Paul-Gilloteaux, P. Roudot, C. Kervrann, F. Waharte, J.-Y. Tinevez, S.L. Shorte, J. Willemse, K. Celler, G.P. van Wezel, H.-W. Dan, Y.-S. Tsai, C. Ortiz de Solórzano, J.-C. Olivo-Marin, and E. Meijering. 2014. Objective comparison of particle tracking methods. Nat. Methods. 11:281–9. doi:10.1038/nmeth.2808.

Daniel, R.A., and J. Errington. 2003. Control of Cell Morphogenesis in Bacteria: Two Distinct Ways to Make Rod Shaped Cell. Cell. 113:767–776. doi:10.1016/S0092-8674(03)00421-5.

Doanachie, K.J.B. and W.D. 1977. Growth of the Escherichia coli Cell Surface. J. Bacteriol. 129:1524.

Errington, J. 2015. Bacterial morphogenesis and the enigmatic MreB helix. Nat Rev Micro. 13:241–248. doi:10.1038/nrmicro3398.

Fishov, I., and C.L. Woldringh. 1999. Visualization of mebrane domains in Escherichia coli. Mol. Microbiol. 32:1166–1172.

Garner, E.C., R. Bernard, W. Wang, X. Zhuang, D.Z. Rudner, and T. Mitchison. 2011. Coupled, circumferential motions of the cell wall synthesis machinery and MreB filaments in B. subtilis. Science. 333:222–225. doi:10.1126/science.1203285.

Joyce, G., K.J. Williams, M. Robb, E. Noens, B. Tizzano, V. Shahrezaei, and B.D. Robertson. 2012. Cell division site placement and asymmetric growth in Mycobacteria. PLoS One. 7:1–8. doi:10.1371/journal.pone.0044582.

Kysela, D.T., P.J. Brown, K. Casey Huang, and Y. V Brun. 2013. Biological Consequences and Advantages of Asymmetric Bacterial Growth. Annu. Rev. Microbiol. 67:417–35. doi: 10.1146/annurev-micro-092412-155622.

Lindner, A.B., R. Madden, A. Demarez, E.J. Stewart, and F. Taddei. 2008. Asymmetric segregation of protein aggregates is associated with cellular aging and rejuvenation. Proc. Natl. Acad. Sci. 105:3076–3081. doi:10.1073/PNAS.0708931105.

Lloyd-Price, J., A. Häkkinen, M. Kandhavelu, I.J. Marques, S. Chowdhury, E. Lihavainen, O. Yli-Harja, and A.S. Ribeiroa. 2012. Asymmetric disposal of individual protein aggregates in Escherichia coli, one aggregate at a time. J. Bacteriol. 194:1747–1752. doi: 10.1128/JB.06500-11.

Nelson, W.J. 2012. Adaptation of core mechanisms to generate cell polarity. Nature. 422:766–774. doi:10.1038/nature01602.Adaptation.

Nielsen, H.J., Y. Li, B. Youngren, F.G. Hansen, and S. Austin. 2006. Progressive segregation of the Escherichia coli chromosome. Mol. Microbiol. 61:383–393. doi:10.1111/j.1365-2958.2006.05245.x.

P. Thévenaz, U.E. Ruttimann, M.U. 1998. A Pyramid Approach to Subpixel Registration Based on Intensity. IEEE Trans. Image Process. 7:27–41. doi:10.1109/83.650848.

Pinho, M.G., and J. Errington. 2003. Dispersed mode of Staphylococcus aureus cell wall synthesis in the absence of the division machinery. Mol. Microbiol. 50:871–881. doi: 10.1046/j.1365-2958.2003.03719.x.

Raskin, D.N., and P.A.J. De Boer. 1999. Rapid pole-to-pole oscillation of a protein required for directing division to the middle of Escherichia coli. Proc. Nat. Acad. Sci. USA. 96:4971–4976.

Rojas, E.R., G. Billings, P.D. Odermatt, G.K. Auer, L. Zhu, A. Miguel, F. Chang, D.B. Weibel, J.A. Theriot, and K.C. Huang. 2018. The outer membrane is an essential load-bearing element in Gram-negative bacteria. Nature. 559:617–621. doi:10.1038/s41586-018-0344-3.

Sbalzarini, I.F., and P. Koumoutsakos. 2005. Feature point tracking and trajectory analysis for video imaging in cell biology. J. Struct. Biol. 151:182–95. doi:10.1016/j.jsb.2005.06.002.

Scheffers, D.J., and M.G. Pinho. 2005. Bacterial cell wall synthesis: new insights from localization studies. Microbiol. Mol. Biol. Rev. 69:585–607. doi:10.1128/MMBR.69.4.585.

Schneider, C.A., W.S. Rasband, and K.W. Eliceiri. 2012. NIH Image to ImageJ: 25 years of image analysis. Nat. Methods. 9:671–675.

Shi, H., B.P. Bratton, Z. Gitai, and K.C. Huang. 2018. How to Build a Bacterial Cell: MreB as the Foreman of E. coli Construction. Cell. 172:1294–1305. doi:10.1016/j.cell.2018.02.050.

Siegal-Gaskins, D., and S. Crosson. 2008. Tightly regulated and heritable division control in single bacterial cells. Biophys. J. 95:2063–2072. doi:10.1529/biophysj.108.128785.

Skerker, J.M., and M.T. Laub. 2004. Cell-cycle progression and the generation of asymmetry in Caulobacter crescentus. Nat. Rev. Microbiol. 2:325–37. doi:10.1038/nrmicro864.

Stewart, E.J., R. Madden, G. Paul, and F. Taddei. 2005. Aging and death in an organism that reproduces by morphologically symmetric division. PLoS Biol. 3:e45. doi:10.1371/journal.pbio.0030045.

Swulius, M.T., and G.J. Jensen. 2012. The helical MreB cytoskeleton in Escherichia coli MC1000/pLE7 is an artifact of the N-Terminal yellow fluorescent protein tag. J. Bacteriol. 194:6382–6. doi:10.1128/JB.00505-12.

Taheri-Araghi, S., S. Bradde, J.T. Sauls, N.S. Hill, P.A. Levin, J. Paulsson, M. Vergassola, and S. Jun. 2014. Cell-Size Control and Homeostasis in Bacteria. Curr. Biol. 25:385–391. doi:10.1016/j.cub.2014.12.009.

Taniguchi, Y., P.J. Choi, G. Li, H. Chen, M. Babu, J. Hearn, A. Emili, and X.S. Xie. 2011. Quntifying E.coli Proteome and Transcriptome with Single-Molecule Sensitivity in Single Cells. Science. 329:533–539. doi:10.1126/science.1188308.

Typas, A., M. Banzhaf, C.A. Gross, and W. Vollmer. 2011. From the regulation of peptidoglycan synthesis to bacterial growth and morphology. Nat. Rev. Microbiol. 10:123. doi: 10.1038/NRMICRO2677.

Ursell, T.S., E.H. Trepagnier, K.C. Huang, and J.A. Theriot. 2012. Analysis of Surface Protein Expression Reveals the Growth Pattern of the Gram-Negative Outer Membrane. PLoS Comput. Biol. 8:e1002680. doi:10.1371/journal.pcbi.1002680.

Varma, A., K.C. Huang, and K.D. Young. 2008. The Min System as a General Cell Geometry Detection Mechanism : Branch Lengths in Y-Shaped Escherichia coli Cells Affect Min Oscillation Patterns and Division Dynamics. J. Bacteriol. 190:2106–2117. doi:10.1128/JB.00720-07.

Varma, A., M.A. de Pedro, and K.D. Young. 2007. FtsZ directs a second mode of peptidoglycan synthesis in Escherichia coli. J. Bacteriol. 189:5692–704. doi:10.1128/JB.00455-07.

Wery, M., C.L. Woldringh, and J. Rouviere-Yaniv. 2001. HU-GFP and DAPI co-localize on the Escherichia coli nucleoid. Biochimie. 83:193–200.

